# Metrics for comparing Neuronal Tree Shapes based on Persistent Homology

**DOI:** 10.1101/087551

**Authors:** Yanjie Li, Giorgio A. Ascoli, Partha Mitra, Yusu Wang

**Affiliations:** Computer Science and Engineering Dept., The Ohio State University, Columbus, OH, U.S.A.; Krasnow Institute for Advanced Study, George Mason University, Fairfax, VA, U.S.A.; Cold Spring Harbor Laboratory, NY, U.S.A.

## Abstract

The geometrical tree structures of axonal and dendritic processes play important roles in determining the architecture and capabilities of neuronal circuitry. Morphological features based on this tree structure have played a central role in classifying neurons for over a century. Yet geometrical trees are not automatically adapted to the basic mathematical tool used widely in data analysis, namely vector spaces and linear algebra, since tree geometries cannot be naturally added and subtracted. Current methods for analysis reduce trees to feature vectors in more or less ad hoc ways. A more natural mathematical object suited to characterizing neuronal tree geometries, is a metric space, where only distances between objects need be defined. In recent years, there have been significant developments in the fields of computational topology and geometry that promise to be useful for the analysis of neuronal geometries. In this paper, we adapt these tools to the problem of characterizing and analyzing neuronal morphology.

As more and more neuroanatomical data are made available through efforts such as NeuroMorpho.org and FlyCircuit.org, the need to develop computational tools to facilitate automatic knowledge discovery from such large datasets becomes more urgent. One fundamental question is how best to compare neuron structures, for instance to organize and classify large collection of neurons. We aim to develop a flexible yet powerful framework to support comparison and classification of large collection of neuron structures efficiently. Specifically we propose to use a topological persistence-based feature vectorization framework. Existing methods to vectorize a neuron (i.e, convert a neuron to a feature vector so as to support efficient comparison and/or searching) typically rely on statistics or summaries of morphometric information, such as the average or maximum local torque angle or partition asymmetry. These simple summaries have limited power in encoding global tree structures. Leveraging recent development in topological data analysis, we vectorize each neuron structure into a simple yet informative summary via the use of topological persistence. In particular, each type of information of interest can be represented as a descriptor function defined on the neuron tree, which is then mapped to a simple persistence-signature. Our framework can encode both local and global tree structure, as well as other information of interest (electrophysiological or dynamical measures), by considering multiple descriptor functions on the neuron. The resulting persistence-based signature is potentially more informative than simple statistical summaries (such as average/mean/max) of morphometric quantities – Indeed, we show that using a certain descriptor function will give a persistence-based signature containing strictly more information than the classical Sholl analysis. At the same time, our framework retains the efficiency associated with treating neurons as points in a simple Euclidean feature space, which would be important for constructing efficient searching or indexing structures over them. We present preliminary experimental results to demonstrate the effectiveness of our persistence-based neuronal feature vectorization framework.

## 1 Introduction

Neuronal cells have a unique geometrical characteristic: tree-like axonal and dendritic processes that can be many orders of magnitude bigger than the cell bodies (somata). These dendritic and axonal trees are fundamental to the operation of neurons, since they enable the coordinated long distance communication of electrical signals, and also enable the complex short and long distance connectivity architecture that is central to nervous system function. In analyzing the circuit properties a data reduction is often made to a connectivity matrix (synaptic connections or mesoscale regional connections), without taking into account the neuronal geometry or topology *per se*. However, it is highly likely that the neuronal geometry plays a critical role in determining the capabilities of the circuit - the geometry is intimately tied to the timing properties of signals in the nervous system and also determine the algorithmic capabilities of the spatially extended circuitry. Since the nervous system enables rapid responses to environmental stimuli to govern behavior, time is of essence. The spatial relations between different inputs to a dendritic tree are important for how the corresponding signals integrate. The tree geometries of neuronal processes reflect developmental dynamics, including the growth and pruning of these processes.

Despite the importance of the geometrical and topological properties of neuronal trees, the characterization and analysis of these properties pose conceptual and methodological challenges. A basic reason for this is the the tree geometries are not naturally characterized by points in some suitable vector space. For tree shapes to be vectors, one should be able to add and subtract tree shapes. There is no natural way to do this. Since vectors spaces (and linear algebra) are fundamental to the data analysis techniques that are widely used, this poses a conundrum. One way out is to map the neuronal geometries to a vector space (through a suitable choice of feature vectors); however, this entails loss of information in a potentially *ad hoc* manner. The alternative is to use a mathematical description that is more naturally suited to tree shapes.

One possibility is to characterize neuronal trees as points in a metric space, where distances between objects are defined, but addition and subtraction of objects need not be defined. While not widely used, metric space techniques do have precedence in neuronal data analysis (eg metric space methods for spike trains). Central to such analyses is a suitable choice of metric or distance between trees. One way to achieve this is to first embed the trees into a vector space, then use a metric in that vector space. However, this intermediate vector space representation could obscure the study of structures that might be present purely in the metric space framework, and also requires the *ad hoc* choice of a vector space representation, so it does not address the basic issue.

In this paper we explore the possibility of directly defining the associated metric space by exploring different tree-metrics, and propose a comprehensive methodology that can also deal with the development or growth of neuron trees and dynamics defined on the trees. The methods may involve reduction to vectors at an intermediate point of analysis, but this happens in a natural and controlled way without ad hoc feature selection. It is also possible to proceed without reduction to vectors, which we indicate but do not pursue in detail in this manuscript. We rely on techniques developed over the last decade based on ideas of topological persistence, that have gained widespread use outside in other applications dealing with geometrical and topological data analysis.

An important application (though not the only one) is to the problem of classifying neurons into classes or types. Axonal or dendritic morphology has been used from early days (cf. Cajal) for such classification purposes, and has been one of the major motivators for past quantitative work based on intermediate feature vector representations. The introduction of computational geometry and topology techniques to this data analysis problem, brings in a modern toolkit, that is also well suited to the large data sets that are becoming available through efforts such as NeuroMorpho.org[4] and FlyCircuit.org[15]. A central question for these data sets is how best to compare neuron structures. This is needed to organize and classify large collections of neurons, to understand variability within a cell type, and to identify features that distinguish neurons. Despite extensive attention from researchers, this problem remains challenging[2]. A broad spectrum of methods to compare neuronal geometries have been developed in this big data context. On one end of the methodological spectrum, the aim is to develop efficient similarity / distance measures for neurons to facilitate efficient classification, search and indexing of neuron data, or as a way to characterize key features of neuron structures. On the other end of the methodological spectrum, the aim is to find a detailed alignment (correspondence) between two or multiple neuron trees to help understand similarity and variation among structures in detail, to help construct consensus or mean structure, and so on. There is typically a trade-off of efficiency versus sensitivity (to structure variation) as we move from one end to the other end of the methodological spectrum.

In this paper, we focus on the efficient end of the spectrum of methods, and aim to develop a flexible yet powerful framework to compare large collection of neuron structures efficiently, while bringing in modern tools for computational geometry and topology.

### Related work

On the efficient end of the method spectrum, there are a family of what one might call *feature-vectorization* methods. Such methods map each neuron structure into a point (a feature vector) in an investigator-defined feature space (often Euclidean space) and the distance between two neurons is measured by the computationally friendly *L*_*p*_-norm between their corresponding feature vectors. Then one can leverage the large literature on searching, nearest-neighbor queries, clustering and classification under *L*_*p*_-norms, to facilitate efficient automatic classification as well as indexing /querying in a big database of neuron structures. One popular way to vectorize a neuron structure is to map it to features consisting of a subset of summarizing morphometric parameters (such as average / max local torque angles) as computed by the L-Measure tool [42]; see e.g, [33, 38, 49]. It has also been observed [2] that classic Sholl-like analysis [43], which counts the number of intersections between neuronal tree with concentric spherical shells centered at soma, provide effective measurements for neuron classification [29, 20, 41]. Other approaches in this family include mapping the skeleton of neuron structure to a density field [44], or representing a neuron by a collection of segments (each represented as a vector) as used in NBLAST [19].

In contrast, on the other end of the method spectrum, at the most sensitive (discriminative) level, one aims to establish (complete or partial) alignments / correspondences between two or multiple neuronal trees, so as to help understand similarity and variation among structures in detail, and to construct a consensus or mean structure. The importance of the specific branching pattern and the tree shape of neurons in their functionality has long been recognized [2]. The neuron structures can be treated as combinatorial trees (where only the connection pattern between nodes matter) or as geometric trees (where locations of nodes and geometric shapes of arcs are also considered). Methods in this category often aim to find correspondences between two (neuron) trees, as well as to develop a tree distance to measure the quality of the resulting tree alignment. One important development in this direction is the use of a tree edit distance (TED) for aligning neuron trees [25, 30]. The tree edit distance can be considered as an extension of the string-edit distance. It measures the distance between two trees by identifying the minimum cost sequence of “edit” operations to convert one tree to the other tree. It is a natural distance for comparing trees, and has been used in various biological applications, such as for comparing phylogenetic trees. Unfortunately, the tree edit distance is NP hard to compute [50], or even to approximate[8]. So current applications use a constrained TED, which can be solved by dynamic programming in polynomial time. The constraints require that ancestor/descendant relations be preserved by the correspondences [25, 30]. The original constrained TED does not model the shape of tree branches, though the alignment used by the multiscale neuron comparison and classification tool BlastNeuron considers the shape of branches to some extent [47]. The DIADEM metric [26] presents a more detailed alignment targeted to the special case of comparing a reconstructed neuron structure with a *gold standard* structure.

In the middle of the spectrum are methods of varying sensitivity and computational costs. Path2Path [5] converts a neuron tree into a set of paths (curves) and then measures distance between two neurons by the distance between corresponding sets of curves. This approach helps to take branch shape into account, but the tree combinatorial structure is somewhat lost. A more enriched model [37] represents a tree as a main curve with several branches (and possibly sub-branches), and uses a dynamic time warping algorithm to align these branches along the main curve. Recognizing the importance of locality of neuronal arborisations, Zhao and Plaza [51] converts neurons into one dimensional distributions of branching density for comparison. Finally, in an interesting recent development [27], Gillette et al. encode the combinatorial structure of a tree as a sequence and compares two or multiple neuron structures using the large literature on sequence alignments.

### New method

In this paper, we focus on the efficient end of the spectrum. We note that current methods to vectorize a neuron typically rely on statistics or summaries of important morphometric information, such as the average or maximum local torque angle or partition asymmetry. These simple summaries have limited power in encoding global tree structures. We develop a new persistence-based feature vectorization framework, which have advantages over previous approaches. First, it provides a unified general framework that can encompass a variety of properties associated with neurons, both static and dynamic. Specifically, each property of interest can be represented as a *descriptor function* defined on the neuron tree, which is then mapped to a simple persistence-signature. This procedure is repeated with other descriptor functions, and the collection of these signatures is considered together as a feature vector. As a result, our framework can encode both local and global tree structure, as well as other information of interest (that pertain to dynamical and electrophysiological properties of neurons), by considering a suitable set of descriptor functions. The resulting persistence-based signature is geometrically meaningful and more informative than simple statistical summaries (such as average/mean/max) of morphometric quantities. As an example, in Section 2.4, we show that by using a natural descriptor function in our framework, our persistence signature is in a mathematically precise sense more informative than the classical Sholl analysis [43]. Secondly, by vectorizing the persistence information in a natural manner, our framework retains the efficiency associated with treating neurons as points in a simple Euclidean feature space. We present some preliminary experimental results to demonstrate the effectiveness of our proposed framework in Section 3. Third, the method generalizes to neuronal shapes that change over time (due to development or experience dependent plasticity), and therefore provides a natural method to capture developmental dynamics.

*Note concering contemporaneous work:* During the course of preparing this manuscript, we were made are of independent work published on the arXiv [31], developed by Kanari et al. Similar to our paper, this paper also proposes to use topological persistence-based profiles to compare neuron morphologies. We would like to note that persistence-based metrics on geometrical graphs have been developed in the prior literature, and do not constitute novel elements in either our work or in the preprint by Kanari et al, but are applications of these literature ideas to neuronal trees. However, our respective applications differ significantly in detail. The work of [31] employs the radial distance function from the root (referred to as Euclidean distance function in our paper). We present a more generic persistence-based feature vectorization framework derived from arbitrary descriptor functions, which has a broader scope. Specifically, as discussed in Section 2.3, we propose to integrate multiple descriptor functions, including functions describing the L-measure, to produce a unified persistence-based signature. Our framework generalizes the Euclidean-distance-based (i.e, radial-distance-based) persistence profile in a similar way that L-measure generalizes the classic Sholl analysis ^1^. We also provide a more detailed roadmap based on our approach suitable for the study of electrophysiological and developmental dynamics. We point out that these two lines of work, despite their similarity, were developed independently. A preliminary presentation of our work was made in poster form at the US BRAIN Initiative annual meeting in December 2015 in Washington DC.

## 2 Persistence-based Signature for Neuron Structures

We model a single neuron as a geometric tree *T* ⊂ ℝ^3^ embedded in the three-dimensional Euclidean space ℝ^3^, where arcs connecting tree nodes are modeled as (polygonal) curves. To incorporate various information of interests on the neuron trees, we model them as *descriptor functions* defined on *T*. We then apply the so-called *topological persistence* to summarize these descriptor functions, to map an input neuron tree (together with various structural or bio-chemical information on it) into a signature (feature vector). The high-level pipeline is shown in Figure 1 – here for simplicity, we use a single descriptor function as an example. But as we describe later, this framework can be extended to multiple descriptor functions.

**Figure 1:**
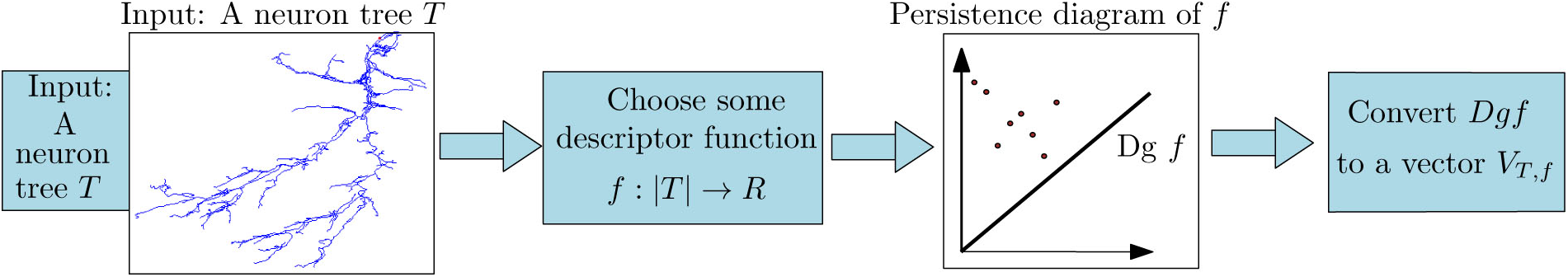
The pipeline of persistence-based feature vectorization framework (for a single descriptor function).

### 2.1 Step 1. Persistence diagram summary

Persistent homology [23] is a basic methodology to characterize and summarize shapes and functions, as well as to identify meaningful features and separate them from “noise” [11, 22]. The underlying space X is examined using a mathematical construct called a *filtration*. A filtration of the space X consists of a nested sequence of indexed subsets of X with the index chosen from an ordered set (such as the set of integers or the set of real numbers), eg X_1_ ⊆ X_2_ ⊆…⊆ X_*n*_ = X. One can think of a filtration to be a specific way to grow and generate *X*. As we “filter” through the space X using this nested sequence of subsets, new topological features may be created and some older ones may be destroyed. Persistent homology tracks the creation (”birth”) and destruction (”death”) of these topological features with respect to the filtration index. For our purposes, we will consider a real valued index, which will call “time”. The resulting births and deaths of features are summarized in a so-called *persistence diagram*. The persistence diagram is set of points in the 2D plane whose (*x, y*) coordinates represent the birth and death times of the features. The life-time of a feature (death time - birth time) is called the *persistence* of this feature, encoding how long this feature exists during the filtration. Since its introduction, persistent homology has become a fundamental method to characterize/summarize shapes, as well as to separate significant features from “noise”.

In our case, given a neuron tree *T*, we first choose one or more real-valued *descriptor functions* defined on *T*. These descriptor functions may encode purely geometric information, such as geodesic or Euclidean distance from a point on the tree, or functions encapsulating electrophysiological or dynamical information (such as the electrotonic distance from a base point). We then use topological persistence to summarize these descriptor functions through suitable filtrations of the neuron tree induced by the descriptor functions.

#### Descriptor functions

We will ignore the thickness of neuronal processes and represent the axonal or dendritic compartment of a neuron as geometric trees *T* embedded in 3D Euclidean space, consisting of tree nodes *V* (*T*) and tree branches (curves connecting the tree nodes). We will use *|T|* to denote the set of points belonging to the tree branches together with the tree nodes. A descriptor function is a real valued function *g*: |*T*| → ℝ defined on |*T*|.

The standard persistence summary that we introduce is defined on descriptor functions defined on a continuous domain. If the function values are specified only at tree nodes *V* (*T*), we can extend these values to a *piecewise-linear (PL)* function *g*: |*T*| → ℝ on |*T*| using linear interpolation along the length the arc. T The main steps involved in the algorithm are shown in Figure 1. In the following, we use the following Euclidean distance descriptor function *f*:|*T*| → ℝ as an example:

Let *r* denote the root of *T*, which may be generically located in the soma of the neuron. Now consider the *Euclidean distance function f*: *T →* ℝ, where for any *x* ∈ *T, f*(*x*) equals the negative of the Euclidean distance between *x* and the root *r* of *T*; that is, *−f* measures how far each point *x* in |*T*| is from the the soma. See Figure 2 (a) for an illustration, where for simplicity we ignore the geometric embedding of the neuron tree *T* and plot it so that the height of each point *x* equals *f*(*x*). Hence we sometimes also refer to *f* as the “height” of a point.

**Figure 2:**
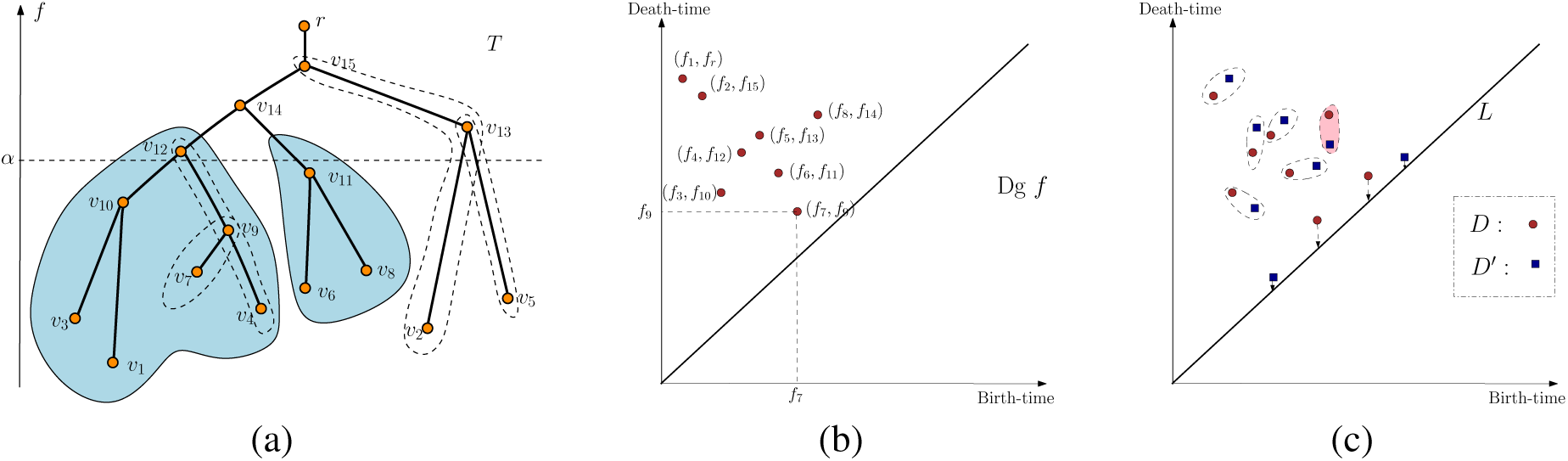
(a) We plot the tree *T* so that the height of a point is its *f* value. The sublevel set *T*_*α*_ is the portion of *T* lying below the horizontal dashed line corresponding to *f* = *α*. Consider what happens when the filtration index *α* passes the vertex *v*_14_. At this time the left and right subtrees (shaded) merge at *v*_14_. These subtrees were originally generated at *m*_1_ = *v*_1_ and *m*_2_ = *v*_6_ respectively. Since the right subtree was born at the later time *f*(*v*_6_) = *f*_6_, this event corresponds to the “death” of the right subtree (with a death time *f*(*v*_14_) = *f*_14_). This gives rise to a persistence point (*f*(*v*_6_), *f*(*v*_14_) = (*f*_6_, *f*_14_) in the persistence diagram in (b). In (b), for simplicity, we set *f*_*i*_:= *f*(*v*_*i*_). We mark some pairs of tree nodes generating persistence points in (a) via dashed closed curves, such as (*v*_7_, *v*_9_) and (*v*_4_, *v*_12_). In (c), red and blue points correspond to persistence points in two persistence diagrams *D* and *D*′, respectively. An example of a correspondence is given, with points matched to diagonal are considered to be noisy points.

#### Persistence diagram w.r.t. *f*

We now describe the persistence diagram summary induced by the so-called *sublevel set filtration* of this function *f*, where *X*_*t*_ = {*x* ∈ *X* |*x < f*(*t*)}. In general, *X*_*t*_ will consist of a set of disjoint pieces of the tree *T*, and birth/death events when a new disjoint piece appears (birth), or two disjointed pieces are joined (a death event for the shorter-lived of the two pieces involved). Note that given the simple topology of the domain *T*, which is a tree, this provides only a simplified view of the general notion of persistent homology.

Consider what happens as we sweep the tree with increasing height values. For any height *α*, we track the connected components in the portion of *T* with height smaller than *α*, called the *sub-level set*

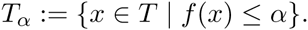

As we sweep past a leaf node, a new component is *created* in the sub-level set. At a saddle point (a branching node), two or more components will be merged into a single one, and thus some components are *destroyed*. Note that each component (a subtree) in the sub-level set *T*_*α*_ is generated (created) by the global minimum in this component (intuitively, this is the first time any point in this component is created). Assume we sweep past a branching point *s* that merges two components, called them *C*_1_ and *C*_2_, into a single component *C*. Suppose *C*_1_ and *C*_2_ are generated by leaf nodes (minima) *m*_1_ and *m*_2_ respectively; and assume w.o.l.g that *f*(*m*_1_) *< f*(*m*_2_). Then intuitively after the merging, the “newer” component *C*_2_ is destroyed and the component *C*_1_ (created earlier at a smaller height) survives with *m*_1_ generating the merged component *C*. As a result, we add a *persistent point* (*f*(*m*_2_), *f*(*s*)) into the persistence diagram Dg*f*, indicating that a feature (branch) originally initiated at height *f*(*m*_2_) is killed at *f*(*s*). The value |*f*(*s*) *− f*(*m*_2_)| is called the *persistence* of this branching feature, specifying its life-time. An example is shown in Figure 2. Two shaded subtrees merge at node *v*_14_, which eliminates the subtree generated at *v*_6_, giving rise to the persistent point (*f*_6_, *f*_14_) in the persistence diagram. Sweeping through the entire tree, we obtain a set of persistent points constituting the persistence diagram Dg*f*, each recording birth and death of branches in a *hierarchical manner* as induced by the distance function *f*. In this paper we use the *extended* persistence diagram, which includes the point (*f*_1_, *f*_*r*_) in the example shown, corresponding to inclusion of the maximum value of the distance function.

Intuitively, we can think of this procedure as a way to decompose the tree into a set of nested branching features (e.g, the feature (*v*_7_, *v*_9_) and (*v*_4_, *v*_12_) in Figure 2), each represented by a point (*b, d*) ∈ Dg*f*, recording its birth and death. Points with larger difference between coordinates have more persistence and so represent more robust features.

#### Super-level sets filtrations

In the above description, we swept the tree bottom up and inspected the changes in components of the sub-level set *T*_*α*_ during the sweep. This procedure captures the “merging” of branching features. Depending on the descriptor function, a general tree could have both down-fork and up-fork nodes (see Figure 3 on the right where we assume that the height represents the function value of tree nodes). Symmetrically, we can also sweep the tree top-down and track the merging in components of the super-level set

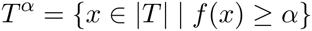

as *α* decreases. This approach would give rise to a set of points recording the splitting-type branching features connected to up-fork tree nodes. We merge the two set of persistence diagrams into a single diagram 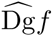 (a single set of planar points), and call it the persistence summary induced by the descriptor function *f*.

**Figure 3:**
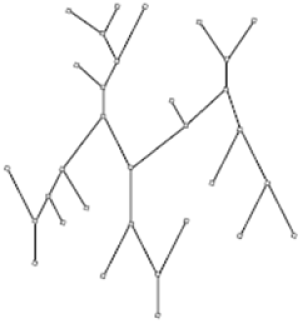
A general tree has both downfork and upfork branching nodes.

Finally, the persistence summary can be efficiently computed in *O*(*n* log *n*) time for an input tree with *n* nodes. Note that this time can be improved to *O*(*n*) time if one assumes that the descriptor function *f* is monotonically increasing along every tree path from root to a tree leaf; see the algorithm used by [31]. However, the *O*(*n*) time complexity does not apply to more general descriptor functions.

### 2.2 Step 2: Vectorization of persistence diagram summaries

Given a neuron tree *T*, we first construct a descriptor function *f* on it, and compute its persistence diagram summary 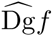 induced by *f*. Given multiple neuron trees *T*_1_,…,*T*_*n*_, we convert each of them to a persistence diagram summary *D*_1_, *D*_2_,…,*D*_*n*_. We now need an efficient way to compute distance between two persistence diagrams so as to compare the corresponding neurons. As we discuss in Appendix A, the standard distance between persistence diagrams used in the topological data analysis literature is the so-called *bottleneck distance* (or its Wasserstein variant [22]). Intuitively, it identifies optimal “almost one-to-one” correspondence between points from one diagramt to the other diagram, so that the maximum distance between pairs of corresponding points is minimized; and this minimal distance is the bottleneck distance between input persistence diagrams. (See Figure 2 (c) for an illustration of a correspondence – some points are allowed to matched to diagonal L:= {(*x, x*)| *x* ∈ ℝ}, in which case they are considered noise.) While is a natural way to measure distance between two persistence diagram summaries, its computation takes *O*(*k*^1.5^ log *k*) where *k* is the total number of persistent points in the diagram. Furthermore, this distance measure does not lend itself easily to fast searching and indexing. Therefore in Step 2, we further vectorize the persistence diagram summaries, to map each persistence diagram into a point in ℝ^*d*^ (i.e, a *d*-dimensional vector) as follows.

Let *D* be a persistence diagram containing points *p*_1_,…,*p*_*k*_ ∈ ℝ^2^. Recall that for each point *p*_*i*_ = (*x*_*i*_, *y*_*i*_), its persistence is |*y*_*i*_ − *x*_*i*_|, which is the vertical distance from *p*_*i*_ to the diagonal *L* = {(*x, x*)| *x* ∈ ℝ}. We can map *p*_*i*_ to a weighted point 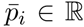 at location *x*_*i*_ with mass |*y*_*i*_ − *x*_*i*_|, which we represent as 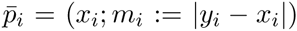. See Figure 4 for an example. Next, we convert the collection of weighted 1D points 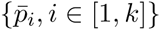 into a 1D density using a simple kernel estimate:

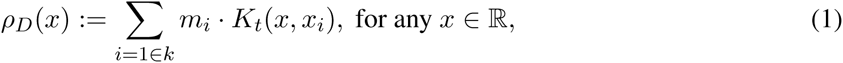

where 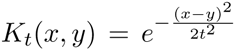 is a Gaussian kernel with width (standard deviation) *t*. We have a Gaussian function *g*_*i*_(*x*) = *m*_*i*_*K*_*t*_(*x*_*i*_, *x*) centered around each *x*_*i*_ and the density function 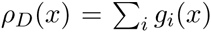 is the sum of these Gaussian functions.

**Figure 4:**
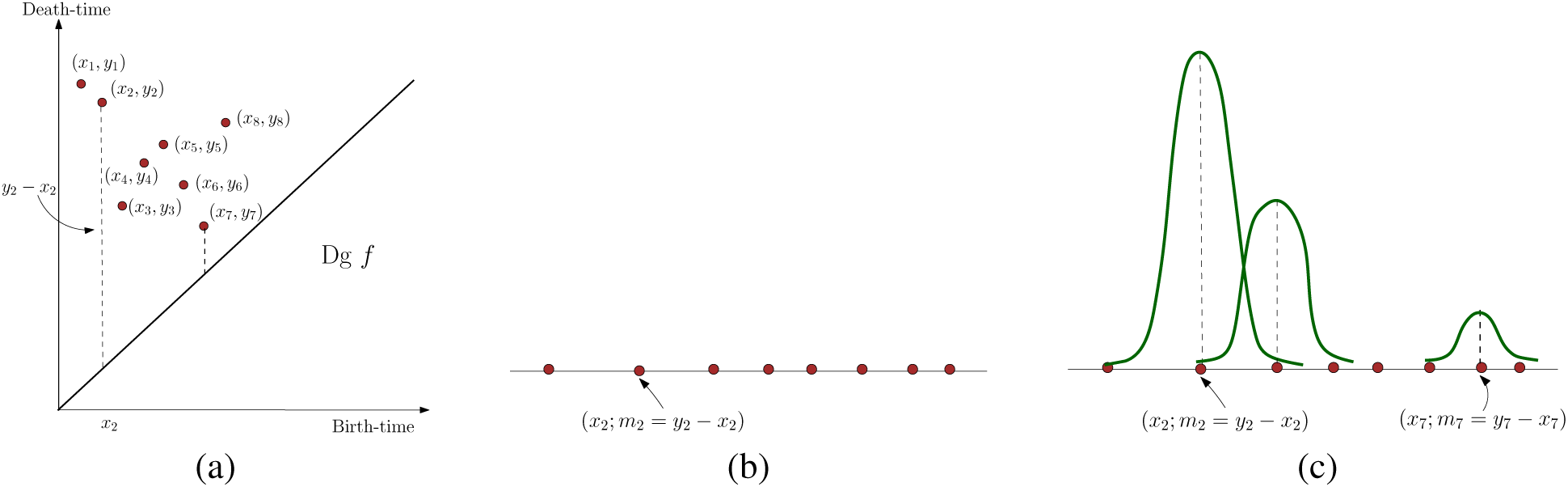
We convert persistent points in the persistent diagram Dg*f* in (a) into a set of weighted points in the line as shown in (b). We then put a Gaussian function *m*_*i*_ · *K*_*t*_(*x*_*i*_, ·) at each point *x*_*i*_ ∈ ℝ, and the sum of them gives the function *ρ*_*D*_. Note that a point with lower persistence (such as (*x*_7_, *y*_7_ *− x*_7_)) has less contribution to the final density function *ρ*_*D*_.

Recall that for a point *p*_*i*_ = (*x*_*i*_, *y*_*i*_) in the persistence diagram, the persistence time |*y*_*i*_ −*x*_*i*_| measures its importance (how long it lives from its birth to death). The weighting of the Gaussian kernel by *m*_*i*_ =|*y*_*i*_−*x*_*i*_| thus gives important features (with larger persistence) greater weights.

Finally, assume that the range of the birth / death times of persistence points in *D* is [a, b] with *I* = b*−*a. We vectorize the density function *ρ*_*D*_ by a *m*-dimensional vector consisting of the function values at the *m* positions evenly spaced in the interval [**a**, b].

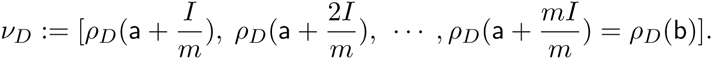

We call the above vector the *persistence-vector*. In our algorithm, we use the same range [a, b] and *m* for all neuron structures, so that their resulting persistent-vectors are comparable.

The distance between two input neurons *T*_1_ and *T*_2_ can then be defined as the standard *L*_*p*_-norm between the resulting vectors 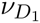 and 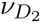 obtained from their persistence profiles *D*_1_ and *D*_2_, respectively. That is, 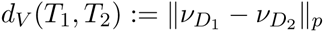.

We remark that there has been several persistence-based profiles developed in the literature of topological data analysis, starting with the *persistence landscape* of [10]. We refer the readers to Section 2 of [1] for a summary of related work. Here we only mention two of the most relevant ones, the multi-scale descriptor of [40] and the persistent images of [1]. Unlike most other persistence-based profiles, both of these two approaches offer some stability guarantees. Our feature vectorization can be considered as a 1D version of the persistent images approach of [1]. We discuss the stability of persistence diagrams and our persistence feature vectors in Appendix A.

### 2.3 Multiple descriptor functions

One advantage of our persistence feature vectorization framework is the generality of the descriptor function *f*. For example, we can use descriptor functions encoding morphometric measurements. Many quantities used in L-measure can induce a descriptor function, such as: (i) define *f*(*v*) to be the branch-angle spanned by the two child-branches of a tree node *v* ∈ *V* (*T*); and (ii) define *f*(*v*) to be the section area or the section radius of the branch at node *v*. We can also consider the *geodesic distance function g*:|*T*| → ℝ, where *g*(*x*) is defined as the geodesic distance to the root of the neuron tree *T*. The descriptor functions can also encode electrophysiological properties. Two such functions are voltage attenuation and propagation delay relative to a base point (e.g. soma), cf. the morphoelectrotonic transform [48].

Furthermore, we can encode more information about an input neuron by using multiple descriptor functions *f*_*i*_s, summarized in the map *F*:|*T*| → ℝ^*r*^:

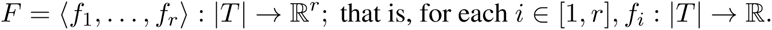

Given *F* = 〈*f*_1_,…,*f*_*r*_〉 defined on *T*, we compute the persistence diagram *D*_1_,…,*D*_*r*_ for each descriptor function. To aggregate these diagram into a single persistent vector, we use the following strategy:

We simply convert each *D*_*i*_ into a feature vector *ν*_*i*_, and concatenate them into a vector *ν*_*T*_ of length *rm*. If the dimension *rm* is too large, later, given a collection of neurons, we perform PCA to reduce the dimension of the resulting persistent-vectors to a lower dimension vector.

#### Potential extensions

As a future work, we will extend the persistence vectorization framework to characterize developmental dynamics of neuronal trees. In particular, the developmental process involves biologically important dynamic changes in neuronal trees. To reflect such changes, the persistence diagram can be extended to handle time-varying data. As a neuron’s structure evolves, the corresponding persistence diagram varies, where each persistence point in it (a branching feature) traces out a curve, called a vine [18]. The evolution of all persistence points traces out a collection of vines, called a vineyard (Figure 5), which summarizes the evolution of a neuron’s structure. Vines can terminate, or new vines can be created, corresponding to the disappearance of an existing branch or creation of a new one. The vectorization procedure can be extended to vineyards, possibly with an intermediate dimensionality-reduction step. Distributed activity measurements (trans-membrane voltage or local calcium concentration) generate a time-varying function that can also be treated in this manner.

**Figure 5:**
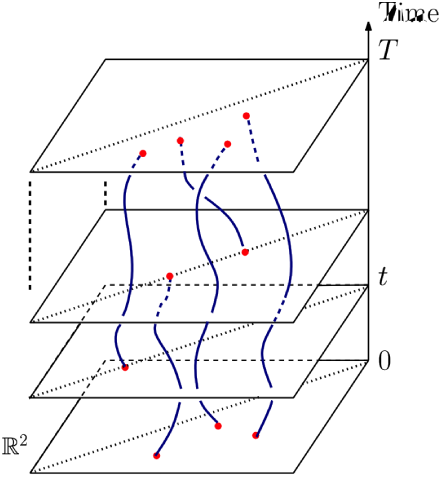
Vertical direction specifies time, and each curve is a vine traced out by a persistent point as time varies.

### 2.4 Connection to Sholl analysis

The persistent-vector for a given descriptor function provides more information than simple statistical summaries such as min, max or average values. Furthermore, by using a geometric descriptor function (such as Euclidean distance function and geodesic distance function), the persistence diagram can encode both local and global shape of the neuron trees, which has been challenging for most previous approaches in comparing neuron trees.

To illustrate the richness of information encoded in persistent summaries, below we show a connection between the persistent diagram Dg*f* of the Euclidean distance function *f* and the previously familiar *Sholl* analysis. Specifically, we show that one can recover quantities used in Sholl analysis from Dg*f*.

Recall that the Sholl analysis is based on the sequence of numbers *N*(*r*) of (dendrite) intersections between a neuron structure and the concentric circle of increasing radius *r* ∈ ℝ^+^, centered typically at the centroid of the cell body. One can treat this count *N* as a function *N*: ℝ^+^ *→* ℝ^+^ w.r.to the radius *r* ∈ ℝ^+^. Various Sholl-type approaches then performs further analysis, such as semi-log analysis of log-log analysis, to obtain one (or more) quantities to summarize this function. Hence the function *N* contains sufficient information for Sholl-type analysis. We call this function the *Sholl function N*.

Next, compute the persistence diagrams Dg^⊥^*f* and Dg^⊤^ *f* induced by the *sublevel set filtration* and the *super-level set filtration* induced by the Eucludean distance function *f*, respectively. Let 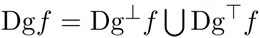 be their union.

Now, consider the *level set f*^−1^1(*r*):= {*x* ∈ |*T*| | *f*(*x*) = *r*} of the function *f*. It can be seen that *N*(*r*) is the number of connected components in the level set *f*^−1^(*r*). As we vary the radius *r* of the concentric circles, components in *f*^−1^(*r*) can appear, disappear, merge and split. The birth and death of components in the level-sets as *r* varies, is recorded by the persistence points in the two persistence-diagrams 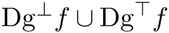 (which is our summary Dg*f*). A persistent point (*b, d*) ∈ Dg*f* indicates that a component is created in the level-set *f^−^*^1^(*r*) with *r* = min{*b, d*}, either as a new component or the splitting of a previous component, and disappears or merges into another component in level-set *f^−^*^1^(*r*′) with *r*′ = max{*b, d*}.

For a connected tree, the value *N*(*r*), for any *r* ∈ ℝ^+^, can be recovend by

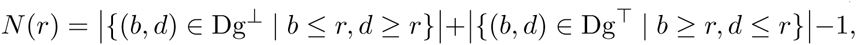

where |*A*| is the cardinality of a set *A*. See Figure 6 for an illustration, where *N*(*r*) is the total number of persistent points in the two shaded quadrants, reduced by one, to account for double counting of the soma or root node. Note that in this paper we utilize the extended persistence diagram (which includes the global maxima/minima of the height function).

**Figure 6:**
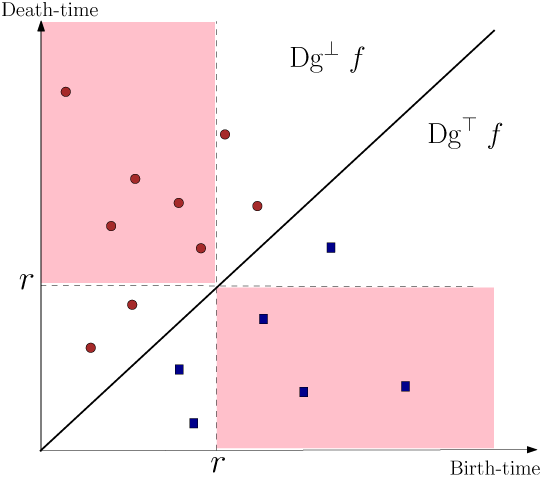
Solid disks are points in Dg^⊥^*f* while squares are points in Dg^⊤^ *f*. *N*(*r*) equals to the number of points in the two shaded quadrants minus one (to correct for double counting of the root node). For the *r* value shown in the picture, *N*(*r*) = 7.

In short, one can retrieve the Sholl function *N* for all *r* values from the persistent summary Dg*f*, and our persistence summary Dg*f* is strictly more informative than the Sholl function *N*. Specifically, while the Sholl function *N* records the number of components in the level set *f^−^*^1^(*r*), the persistent summary Dg*f tracks* these components – Indeed, as mentioned earlier and recall Figure 2, the persistent homology intuitively produces a hierarhical family of nested *branching features*, and each point in the persistence diagram encodes one such feature.

## 3 Preliminary Experimental Results

In this section, we provide some preliminary experimental results to as a proof-of-principle demonstration of our framework. In particular, we show the effectiveness of our method with just a single descriptor function. The descriptor function we choose is the Euclidean distance function: This natural descriptor function encodes information from Sholl Analysis as explained in Section 2.4, and the persistence induced by this function reflects the morphology of the tree underlying an input neuron.

We use three test data sets below. The first test data set Dataset 1 is taken from [45], consists 379 neurons, taken from the Chklovskii archive (Drosophila) of NeuroMorpho.Org, manually categorized into 89 types [51]. All the skeletons including the type information can be downloaded from http://neuromorpho.org under the Drosophila - Chklovskii category.

The second test dataset Dataset 2 contains 114 neurons from four families: Purkinje, olivocerebellar neurons, Spinal motoneurons and hippocampal interneurons, downloaded also from neuronmorpho.org. Specifically, the 16 Purkinje reconstructions [39, 46, 14, 35] have only dendrites with no axons. The 68 olivocerebellar neurons reconstructions [9] have only axons, with no dendrites. The 17 spinal motoneurons reconstructions [32] have complete dendrites, but only the initial branches of the axons. In this case, we keep only the dendrites in our experiments. The 13 hippocampal interneurons reconstructions [3] have both dendrites and axons. In this case, we separate each hippocampal interneuron reconstruction into two trees: one for dendrites and one for axons. In total, we obtain 127 neuron trees, some of them are dendrite trees and some are axonal trees.

The third test data set Dataset 3 comes from the Human Brain Project [34] and downloaded from NeuroMorpho.Org. It includes 1268 neuron cells, out of which the primary cell class is known for 1130 cells, and there are two primary cell classes: interneurons and principal cells. Both of these classes have complete dendrites. The interneurons have moderately complete axons, the principal cells have incomplete axons. We have not separated the dendrite and axonal trees in this case.

### Nearest-neighbor classification accuracy

In this test, we aim to demonstrate the discriminative power of the persistence profiles. Given an input set of neurons 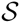, we compute the persistence diagram *D*_*T*_ for each of the input neuron 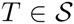, and represent 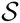 by 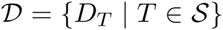. We further vectorize these persistence diagrams and obtain a collection of feature vectors 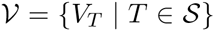. To understand the effect of feature vectorization, below we will consider two distance metrics between neurons:

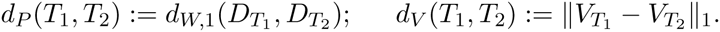

That is, *d*_*P*_ is a distance based on the persistence diagram representation, defined as the degree-1 Wasserstein distance between the two persistence diagrams 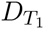 and 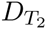 (see Eqn (3) in Appendix A for the definition of 1-Wasserstein distance). *d*_*V*_ is the *L*_1_-distance based on the persistence feature vector representation.

We next perform simple nearest-neighbor classification to decide the class membership of a query neuron: That is, given a query neuron *T*, we compute its nearest neighbor *T*′ in the training set under either *d*_*P*_ or *d*_*V*_, and return the class membership of *T*′ as that for *T*. To test the accuracy of this simple classification, we perform the leave-one-out cross validation. Specifically, for each neuron 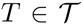, we take 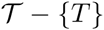 as the training set, and find its nearest neighbor in 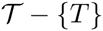. We consider it a *success* if its nearest neighbor is from the same cell-type class as *T*.

We further extend this to hit-rate for the top *k*-nearest neighbor: that is, given a neuron 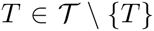, we compute its *k*-nearest neighbors in 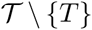, and consider it a “success” if these *k*-NNs include a neuron from the same family of *T*.

The results are reported in Table 1. For Dataset 1, we consider only those classes with at least 2 members, as otherwise, it is meaningless to classify a neuron *T* when 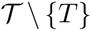 does not contain any neurons from the same family as *T*. This leaves us 346 number of neurons. For Dataset 3, we consider only those neurons whose class-memberships are known, which gives us 1130 cells.

**Table 1:**
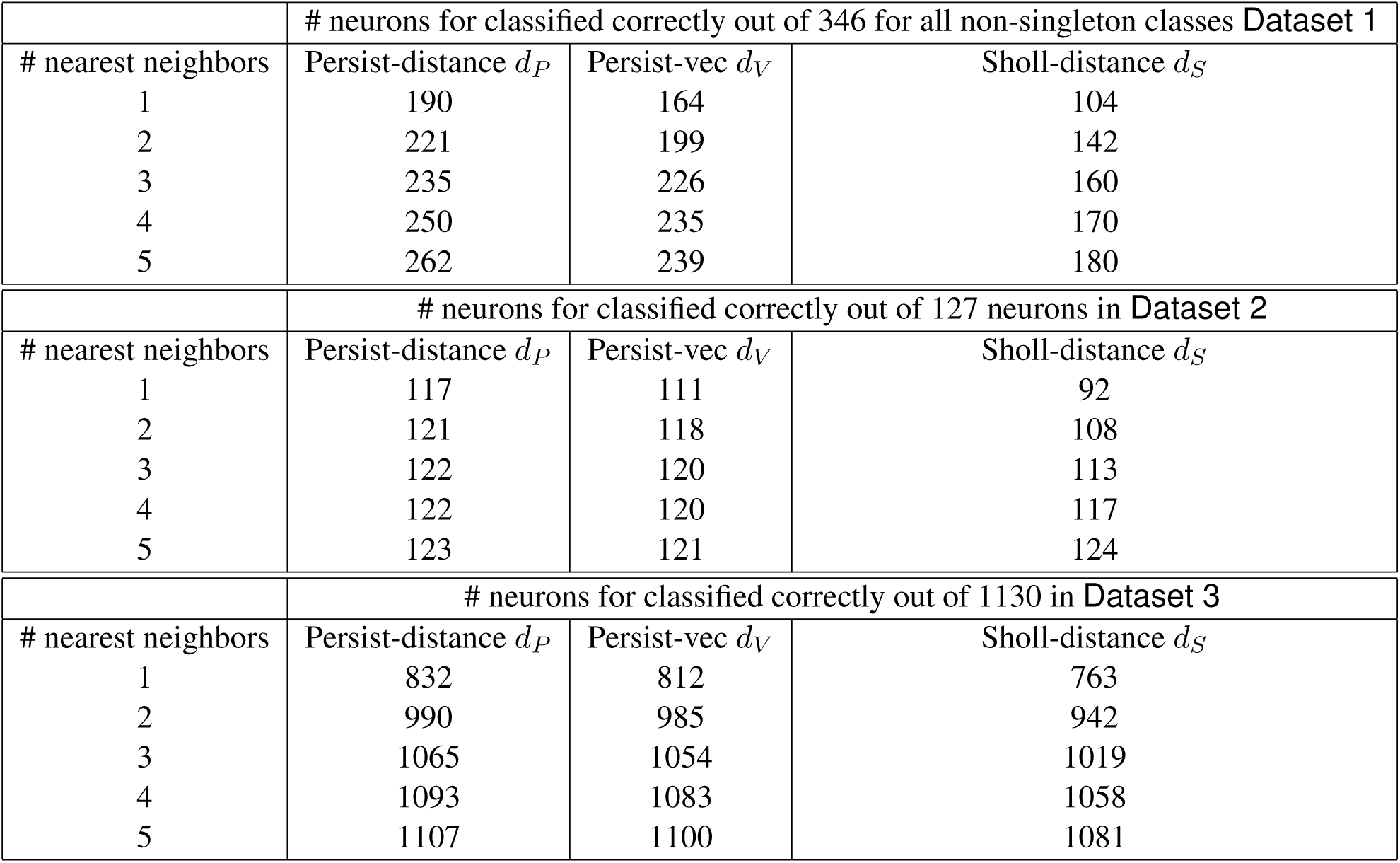
Leave-one-out cross validation for *k*-nearest neighbor classification rate where *k* = 1, 2,…,5. For Dataset 1, we only keep classes which contains at least 2 members.

In the table, each column shows the number of successes (that is, how many time the nearest neighbor of the query is from the same class of it) under three different distance metrics between neurons. Column 2 is based on the Wasserstein distance between persistence diagrams *d*_*P*_, and Column 3 shows results based on the *L*_1_-distance between vectorized persistence diagrams *d*_*V*_. For comparison purposes, in Column 4, we use a distance *d*_*S*_ based on Sholl-type analysis. In particular, given an embedded neuron structure *T*, we compute the Sholl function *N*_*T*_: ℝ^+^ *→* ℝ^+^ as introduced in Section 2.4, where *N*_*T*_ (*r*) equals the number of intersections between the neuron tree *T* and the radius-*r* sphere centered at the root of tree *T*. Given two neurons *T* and *T*′, intuitively, we would like to define the Sholl-based distance *d*_*S*_(*T, T*′) as the *L*_1_-norm of of *N*_*T*_ − *N*_*T*_. In our implementation, we discretize each Sholl function to a vector 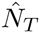 of size 100 (which is the same as the size of discretization for the persistence feature vector), and compute the *L*_1-_ distance between the two vectors as 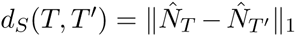. We note that *d*_*S*_ directly compare the Sholl functions (profiles) and thus tends to be more discriminative than using summary quantities, such as the area below the Sholl functions, or semi-log / log-log Sholl analysis, often used to compare neuron morphologies.

We note that using persistence diagrams give slightly better results than using persistence vectors; but we do not lose much discriminative power by using the persistence vectors. We also note that using the Sholl function gives worst classification accuracy, especially when the number of nearest neighbors is small. The difference is particularly prominent for Dataset 1, which is more diverse than other datasets. This indicates that our persistent-based summary indeed is more discriminative than simple statistical summary even though they come from the same descriptor function.

We remark that the branch-density-based similarity measure proposed in [51] gives better classification accuracy over Dataset 1. However, we note that using the Euclidean distance descriptor function, our approach is rigid-transformation invariant. The method of [51] however assumes that the input neurons are from the same coordinate system (with column / tangential directions given).

### Clustering

We now explore the clustering structure of input neurons based on our persistence-based distance. In Figure 7 (a), we show the embedding of the 127 neurons in Dataset 2 to the plane via Laplacian Eigenmap [7], which is a popular non-linear dimensionality reduction method. Each node in the plot represents a neuron, and its color reflects its neuron type. As we can see from the plot, neurons of different types are separated. To understand the clustering structure in more detail, we perform the so-called average linkage clustering method to produce a hierarchical clustering (HC) of the input neurons. In a hierarchical clustering tree (HCT), each leaf corresponds to an input neuron, each subtree represents a cluster, a down-fork node indicates the merging of two or more clusters (subtrees), and its height value corresponds to at which distance threshold this merging happens. The HCT of Dataset 2 is shown in Figure 7 (b), where leaf nodes (neurons) are marked by the same color coding as in (a).

**Figure 7:**
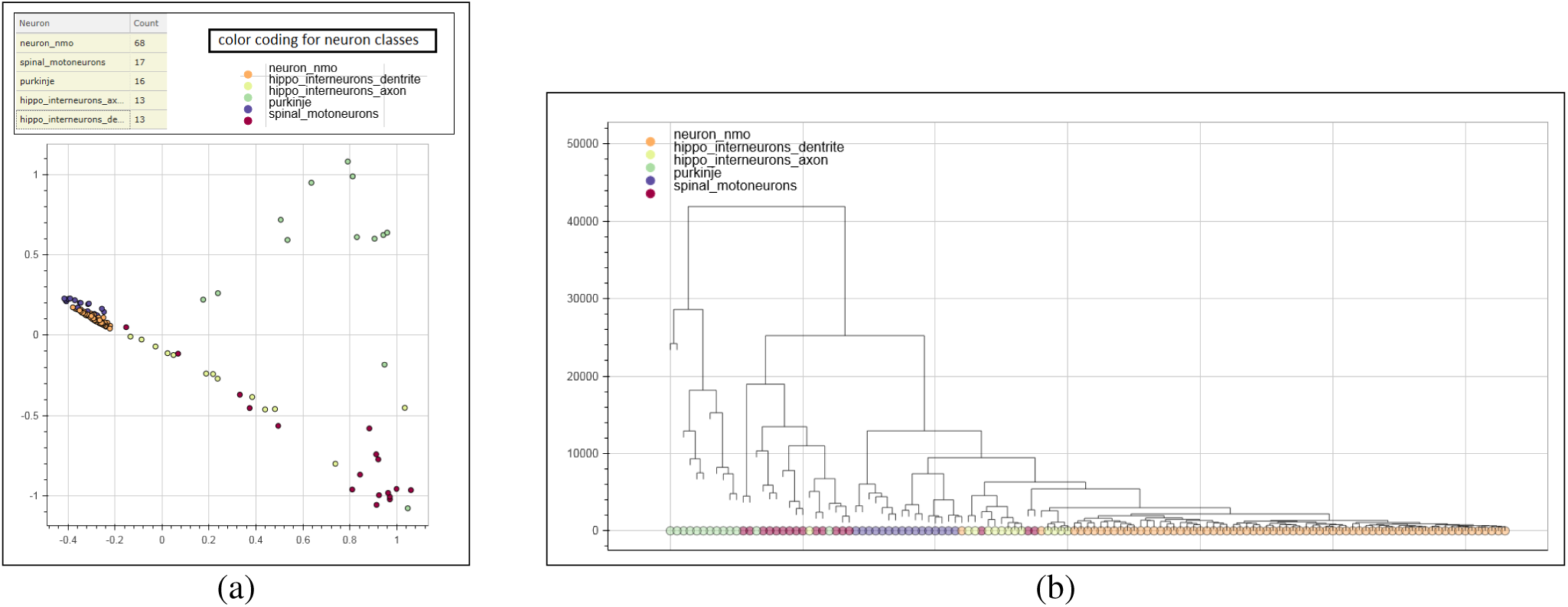
(a) Embedding of Dataset 2 in 2D via Laplacian Eigenmap. Each dot represents a neuron, and its color corresponds to its type. Its hierarchical clustering tree (HCT) is shown in (b), where each leaf corresponds to a neuron.

We choose a hierarchical clustering method since (1) it reveals more sub-clustering information than a flat clustering method, and (2) it permits the construction of a visualization platform based on the hierarchical clustering structure to allow users to interactively explore the input data.

In Figure 8, we show the hierarchical clustering tree (HCT) for Dataset 1. Each leaf node corresponds to a neuron, and we mark those from top 5 largest classes of the 89 manually categorized classes [45] by colors – The largest class “Tangential” is excluded in this figure: Members from this class spread into several clusters in the HCT. This class is proved challenging to classify in the previous work as well [51]. As we can see, majority of neurons from each class are clustered together in the hierarchical clustering tree.

**Figure 8:**
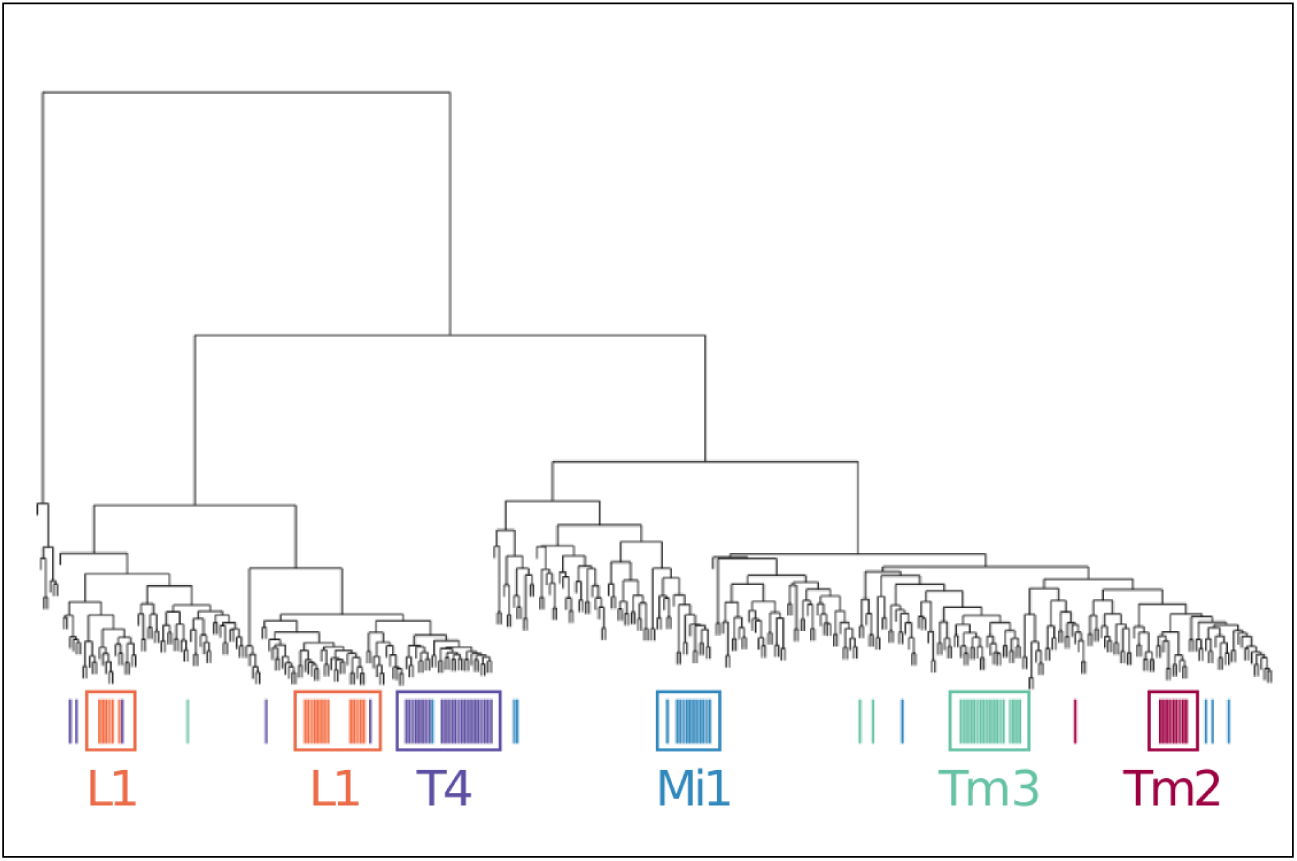
The HCT of Dataset 1. Each leaf corresponds to a neuron structure, and those from the top 5 largest classes (class “Tangential” excluded) are marked (color coded).

### Visualization of the space of neurons

While the visualization of HCT in Figures 7 and 8 is useful in studying the clustering structure behind a collection of neurons, such a tree-visualization becomes ineffective for large data sets, mainly due to the cluttering of the large number of nodes. Indeed, the HCT already becomes hard to interact for our Dataset 3 with only a little more than 1000 structures. At the same time, the number of available neuron structures is rapidly increasing. For example, NeuroMorpho.org holds about 50,000 structures just within a few years of its establishment.

We have developed a terrain visualization for the HCT, based on the Denali framework [24]. Specifically, instead of showing a tree, we build a terrain in 3D corresponding to the input HCT. Each peak of the terrain corresponds to a cluster (i.e, the collection of nodes within some subtree in the HCT), and when two clusters merge in the HCT, their corresponding peak merges in the terrain. This terrain visualization platform provides many functionalities (see [24] for details), including allowing the coloring of the terrain based on a property of interest. A very important functionality is that the platform allows the user to explore a selected group of neurons in details, as well as to inspect each individual neuron structure. In particular, when a user clicks a specific region (corresponding to one cluster which main contain multiple levels of sub-clusters), our tool will return all the neurons contained in that cluster, and plot (i) the subtree rooted at this node, which corresponds to the multiple-levels of subclusters contained in this cluster and (ii) an embedding of all neurons in this cluster to the 2D plane via Laplacian Eigenmap. Note that these two types of visualization are not effective for large data sets, but effective now for a single cluster, which is typically of much smaller size. Furthermore, the user can select each individual neuron from either plots, and when a neuron is selected, its corresponding geometric structure will be shown in another panel which allows interactive manipulation. Other accompanying information (such as L-measure values) will also be shown if available. Thus we have developed a tool for exploring the abstract space of neuronal morphologies using our persistence-based distance.

### Support

This work was supported by the Crick-Clay Professorship at CSHL to Dr Mitra, NSF BRAIN Eager 1450957 and NIH 1R01EB022899-01.

**Figure 9:**
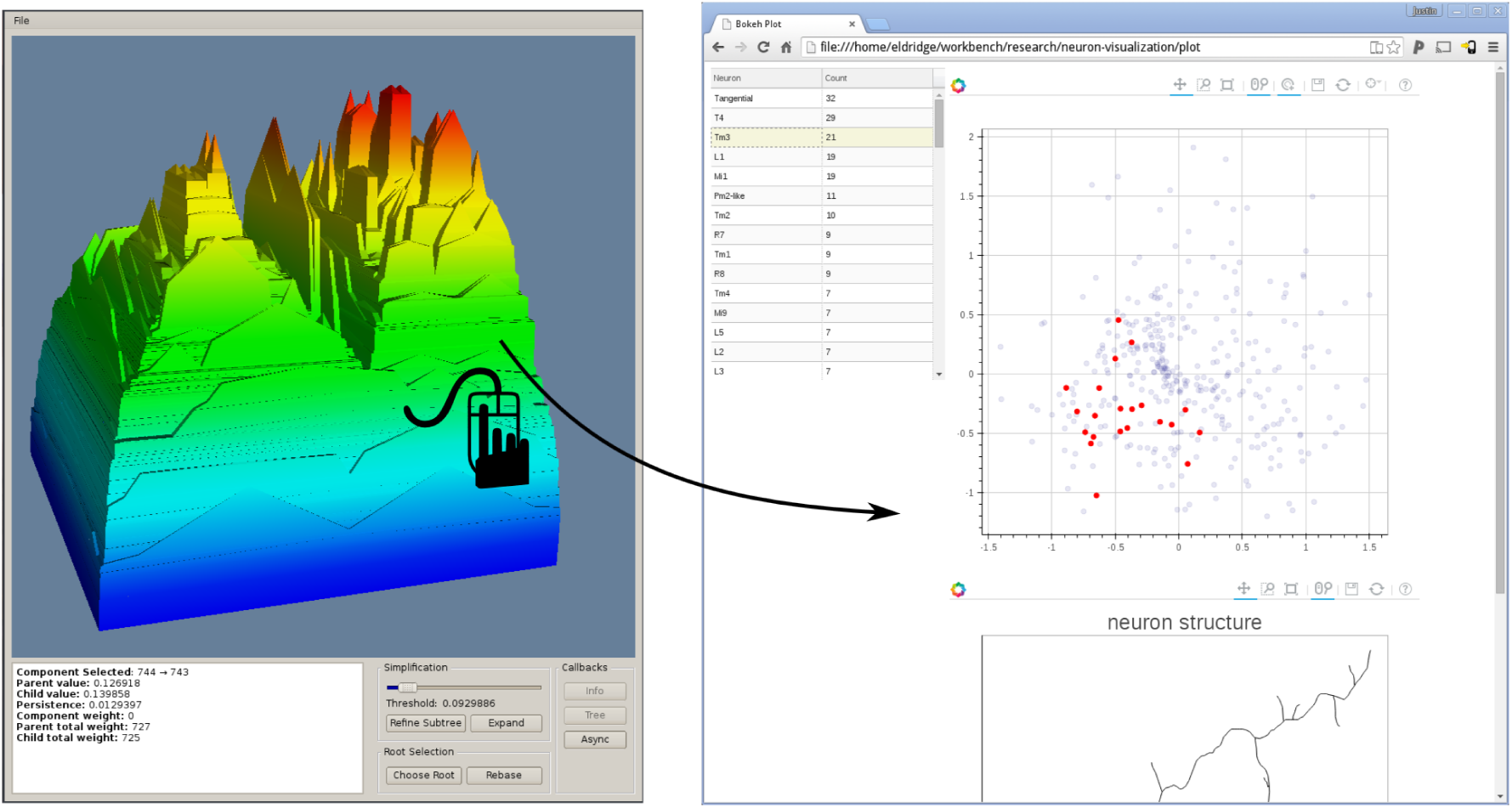
As a user selects a region in the terrain, the HCT view and embedding view are shown on the right. One can further select a neuron from the HCT / 2D embedding view, and inspects its structure as well as associated L-measure information.

## A Stability of persistence-based signatures

Given two neuron trees *T*_1_ and *T*_2_, in (Step 1) we first map them to their respective persistence diagrams *D*_1_ and *D*_2_ induced by some descriptor function(s). In (Step 2), we further vectorize these persistence diagrams into persistence feature vectors, say *V*_1_ and *V*_2_ respectively. It is desirable that such a feature generation and vectorization process is *stable* in the sense that “small perturbations” in input neuronal trees and in the induced descriptor functions should only cause small changes in the distances between them. Making such a stability statement precise is not trivial, depending also on how “perturbations” are modeled and measured. While we do not yet have a full stability statement for our persistence-based feature vectors, below we discuss some partial results. We separately consider the stability for persistence diagrams (after Step-1) and that of persistence-based feature vectors (Step-2).

### Stability of persistence diagrams

To discuss stability, we first need to measure the distance between two persistence diagrams. Given two persistence diagrams *D*_1_ and *D*_2_ (each of which consists of a set of points in IR^2^), there is a natural distance measure, the *bottleneck distance d*_*B*_(*D*_1_, *D*_2_) first introduced in [16]. Consider matching points in *D*_1_ with points in *D*_2_ such that each point in *D*_1_ (resp. in *D*_2_) has to be matched, either to a unique point in *D*_2_ (resp. in *D*_1_), or to its nearest neighbor in the diagonal L:= {(*x, x*)| *x* ∈ ℝ}: The latter case corresponds to treating this feature point as noise, in which case it is matched to a persistent point with zero persistence. See Figure 2 (c) for an illustration. Find the optimal correspondence so that the maximum distance between pairs of corresponding points is minimized; *d*_*B*_(*D*_1_, *D*_2_) equals this smallest possible maximum distance.

More precisely,

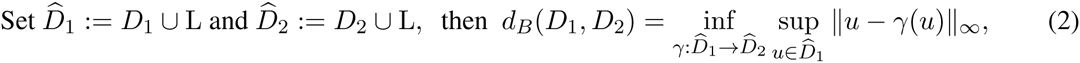

where *‖u − v‖_∞_* = max{*|u.x − v.x |,|u.y − v.y |*} denotes the *L*_*∞*_ distance between two points; and *γ* ranges over all bijections between 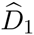 and 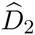. It is known that the bottleneck distance *d*_*B*_(*D*_1_, *D*_2_) can be computed in *O*(*k*^1.5^ log *k*) time, where *m* is the total number of points in *D*_1_ ∪ *D*_2_.

Given two functions *f, g*:|*T*| → ℝ, suppose that *g* is a perturbation of *f* with bounded distance in *L*_*∞*_-norm, that is *‖f − g‖_∞_*:= max_*x*∈|*T*|_|*f*(*x*) *− g*(*x*)| measures the amount of perturbation of *g* from *f*. The Stability Theorem [16] states that for a function *f* and its perturbation *g*, the bottleneck distance between their persistence diagram summaries is bounded from above by the size of the perturbation; that is, *d*_*B*_(Dg*f*, Dg *g*) ≤ ‖*f − g*‖_∞_. This result is later generalized to more general persistence modules, and show that the bottleneck distance between two persistent diagrams is bounded by the so-called interleaving distance between the corresponding persistence modules that generate them [12, 13].

In our setting, given two neuron trees *T*_1_ and *T*_2_ with descriptor functions *f*_1_:|*T*_1_| *→* ℝ and *f*_2_:|*T*_2_| *→* ℝ, we cannot directly compare these two descriptor functions since they are defined on different domains (*T*_1_ and *T*_2_, respectively). We instead use the so-called *functional distortion distance* [6] *d*_*F D*_(*f*_1_, *f*_2_) to measure how different the functions *f*_1_ and *f*_2_ are. Intuitively, *d*_*F D*_ considers all pairs of mappings between *T*_1_ and *T*_2_ as a way to align them, say *ϕ*:|*T*_1_| *→ |T*_2_| and *ψ*:|*T*_2_| *→ |T*_1_| to align *T*_1_ to *T*_2_ (via *ϕ*) as well as align *T*_2_ to *T*_1_ (via *ψ*). It then compares *f*_1_ and *f*_2_ composited with these maps so that they are then defined on a common domain. Each such pair of maps (alignment) (*ϕ, ψ*) will give a cost, measuring how well *f*_1_ and *f*_2_ are aligned under these two maps, and *d*_*F D*_(*f*_1_, *f*_2_) returns the minimum cost under all possible such alignments (pairs of maps). We refer the readers to [6] see the formal definition. It follows from results of [6] that *d*_*B*_(Dg*f*_1_, Dg*f*_2_) ≤ *d*_*F D*_(*f*_1_, *f*_2_).

This stability result applies to any descriptor functions. For example, suppose we consider the Euclidean distance functions *f*_1_ and *f*_2_ on the two trees in Figure 10. Then their persistence diagram summaries are close (at most *ε*), despite that there are noisy branches, as well as combinatorial changes in the tree structures from *T*_1_ to *T*_2_ – Indeed, it is not hard to establish maps *ϕ*: *T*_1_ *→ T*_2_ and *ψ*: *T*_2_ *→ T*_1_ and show that the cost of the Euclidean distance function incurred by them is at most *ε*, thus upper bounds *d*_*F D*_(*T*_1_, *T*_2_) by *ε* too.

**Figure 10:**
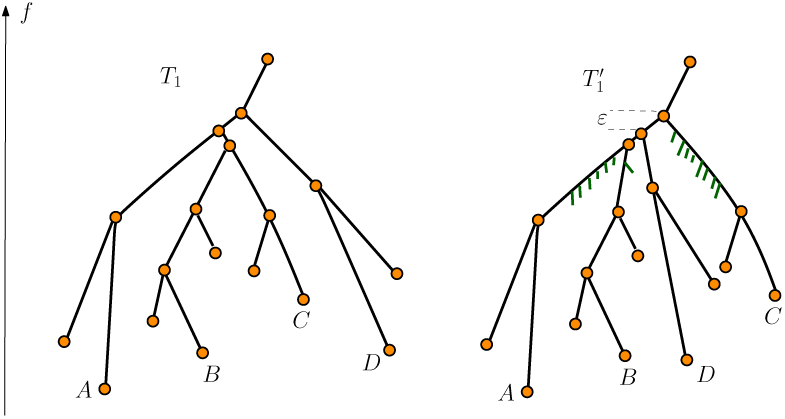
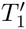 is a noisy version of *T*_1_: there could be local combinatorial changes such as the subtrees *B, C* and *D* merge at slightly different height, and there could also be spurious noisy branches. However, such changes do not perturb the tree metric much: in this specific example, the metric distortion is bounded by *ε*. As a result, their corresponding persistence diagram summaries are also close with *d*_*B*_(*D, D*′) *≤ ε*.

As another example, if we use the geodesic distance to the root as the descriptor function, then by using results from [21], the bottleneck distance between resulting persistence diagrams is stable w.r.t. changes in the input neuron trees as measured by the Gromov-Hausdorff distance between these trees. The Gromov-Hausdorff distance is popular way to measure the level of near-isometry between two metric spaces [28, 36]. The Gromove-Hausdorff distance between the two neuron trees in Figure 10 is at most *ε*, implying that using the geodesic distance as descriptor functions *f*: *T*_1_ *→* IR and 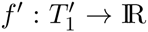, we have *d*_*B*_(Dg*f*, Dg*f*′) *≤ ε* as well.

In our framework, to improve computational efficiency, we vectorize the persistent diagrams describe in (Step 2), and the natural *L*_*p*_-distance between them are sum-based, instead of max-based (as in bottleneck distance). One can extend the bottleneck distance to the so-called degree *p*-Wasserstein distance *d*_*W,p*_(*D*_1_, *D*_2_) between two persistence diagrams *D*_1_ and *D*_2_ which we will introduce shortly in Eqn (3). The stability for the Wasserstein distance of persistence diagrams is not as well understood as in the bottleneck distance case (which is in fact the case when *p* = *∞*), although there are some results for some special cases [17].

### Stability of the persistence feature vectors

We now discuss the stability of feature vectorization step. Specifically, suppose we are given two persistence diagrams *D*_1_ and *D*_2_, with corresponding feature functions 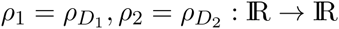 IR induced from *D*_1_ and *D*_2_ as described in Section 2.2.

First, the *degree-p Wasserstein distance between D*_1_ *and D*_2_, for 1 *≤ p < ∞*, is defined as:

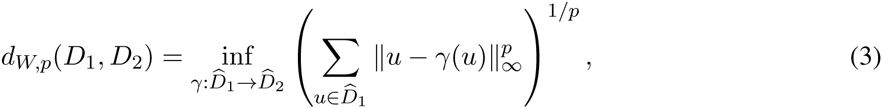

where 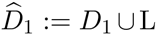 and 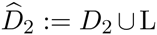 are the persistence diagrams augmented with points in the diagonal L as before.

#### Theorem A.1

*The L*_1_-*distance between feature vectors V*_1_ *and V*_2_ *is stable w.r.to the* 1*-Wasserstein distance between the diagrams D*_1_ *and D*_2_ *generating them. Let* Δ *denote the largest persistence value of any point in D*_1_*; that is*, 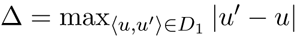. *Specifically*,

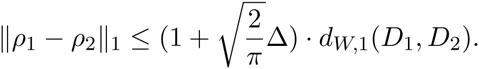

We now prove this theorem. First, we need the following result bounding the distance between two 1-dimensional Gaussians [1].

#### Lemma A.2

([1]) *Given u* ∈ IR, *let g*_*u*_: IR *→* IR *denote the normalized 1-dimensional Gaussian centered at* 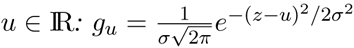. *For a, b* > 0 *and u, v* ∈ IR, *we then have*:

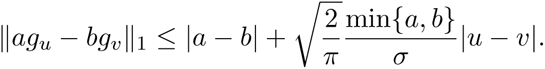

Now let 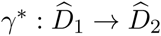 denote the optimal bijection to give rise to *d*_*W*,1_(*D*_1_, *D*_2_); that is,

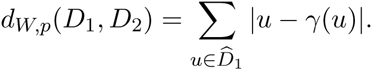

Recall by Eqn 1, for *i* = 1 or 2,

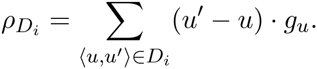

Since for all points (*u, u*′) ∈ L in the diagonal L, we have that *u* = *u*′. It then follows that equivalently, we have that

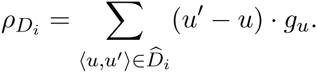

Finally, for each point 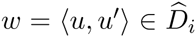, we also denote *w.birth* = *u* (the birth time of this point) and *w.pers* =|*u′ − u*| (the persistence of this point). We now have:

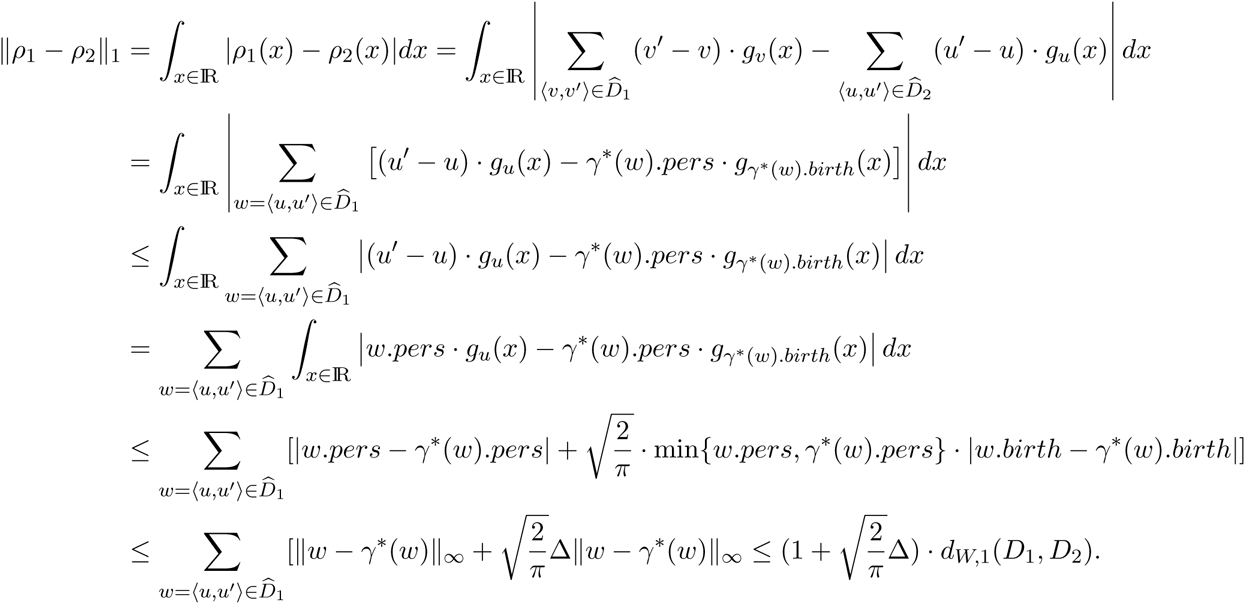

Theorem A.1 then follows.

